# Multiple lesion inductions intensify central sensitization driven by neuroinflammation in a mouse model of endometriosis

**DOI:** 10.1101/2025.01.23.634555

**Authors:** Madeleine E. Harvey, Mingxin Shi, Yeongseok Oh, Debra A. Mitchell, Ov D. Slayden, James A. MacLean, Kanako Hayashi

## Abstract

**Introduction:** Endometriosis is an inflammatory disease associated with chronic pelvic pain (CPP). Growing evidence indicates that endometriotic lesions are not the sole source of pain. Instead, central nervous system (CNS) dysfunction created by prolonged peripheral and central sensitization plays a role in developing endometriosis-associated CPP. This study investigated how CPP is established using a multiple lesion induction mouse model of endometriosis, as repeated retrograde menstruation is considered underlying endometriosis pathogenesis.

**Methods:** We generated endometriosis-like lesions by injecting endometrial tissue fragments into the peritoneal cavity in mice. The mice received a single (1x) or multiple inductions (6x) to simulate recurrent retrograde menstruation. Lesion development, hyperalgesia by behavioral testing, signs of peripheral sensitization, chronic inflammation, and neuroinflammation were examined with lesions, peritoneal fluids, dorsal root ganglia (DRG), spinal codes, and brain.

**Results:** Multiple lesion inductions increased lesion numbers and elevated abdominal and hind paw hypersensitivity compared to single induction mice. Elevated persistent glial cell activation across several brain regions and/or spinal cords was found in the multiple induction mice.

Specifically, IBA1+ microglial soma size was increased in the hippocampus and thalamus. IBA1+ cells were abundant in the cortex, hippocampus, thalamus, and hypothalamus of the multiple induction mice. GFAP+ astrocytes were mainly elevated in the hippocampus. Elevated TRPV1, SP, and CGRP expressions in the DRG were persistent in the multiple induction mice. Furthermore, multiple inductions induced the severe disappearance of TIM4^hi^ MHCII^lo^ residential macrophages and the influx of increased proinflammatory TIM4^lo^ MHCII^hi^ macrophages in the peritoneal cavity. The single and multiple inductions elevated secreted TNFα, IL-1β, and IL-6 levels in the peritoneal cavity at 2 weeks. Elevated cytokine levels returned to the pre-induction levels in the single induction mice at 6 weeks; however, they remained elevated in the multiple induction mice.

**Conclusions:** Our results indicate that the repeatedly occurring lesion inductions (=mimic retrograde menstruation) can be a peripheral stimulus that induces nociceptive pain and creates composite chronic inflammatory stimuli to cause neuroinflammation and sensitize the CNS. The circuits of neuroplasticity and stimulation of peripheral organs via a feedback loop of neuroinflammation may mediate widespread endometriosis-associated CPP.

## Introduction

Endometriosis is a chronic inflammatory disease characterized by the presence of endometrium-like tissues outside the uterus [1] that affects approximately 10% of reproductive-aged women, representing ∼190 million women worldwide [2, 3]. It can cause debilitating chronic pelvic pain (CPP), manifesting dysmenorrhea, dyschezia, dysuria, dyspareunia, and acyclic pelvis pain that dramatically reduces the quality of life of women [4–7]. Many patients can endure symptoms for several decades due to the onset of endometriosis-associated pain during adolescence [3] and have a greater risk of chronic opioid use for pain relief [8]. Despite a sizeable clinical burden, the pathogenesis of endometriosis is complicated and remains poorly understood. The current medical treatment/management is non-curative. It is limited to surgical excision of endometriotic lesions and/or hormonal treatments to suppress estrogen production and action due to endometriosis being an estrogen-dependent disease. Surgical excision of lesions can alleviate endometriosis-associated pain, though pelvic pain frequently returns within a year of lesion removal, even in the absence of lesion regeneration [9, 10]. Thus, endometriosis-associated CPP is not solely dependent on the presence of lesions [11].

Pain relies on peripheral stimuli to the spinal cord for processing and perception by the brain. Inflammatory mediators, such as proinflammatory cytokines and chemokines, prostaglandins, and NGF, evoke pain by directly activating and sensitizing nociceptor neurons in the peripheral tissues via modulation of various ion channels like TRPA1, TRPV1, and voltage-gated sodium channels [12]. Sensitized and activated nociceptors, specifically C-fibers, secrete neuropeptides like SP and CGRP [13], which can trigger a positive feedback loop to stimulate proinflammatory mediator secretion, further perpetuating pain signaling [11]. Through these processes of sensory signal transduction, increased release of neurotransmitters, such as SP and CGRP, induces hyperactivity and hypersensitivity in the spinal cord and brain, known as central sensitization [14]. In endometriosis, abundant immune responses are present at lesion sites with increased proinflammatory cytokines and chemokines, growth factors, and NGF found throughout the pelvic cavity [15–18]. Elevated TNFα, IL-1β, and IL-6 levels have been reported in the peritoneal fluids and/or eutopic and ectopic endometrial tissues of women with endometriosis [17, 19–21]. Specifically, TNFα, IL-1β, CLL5, and NGF are elevated in the pelvic cavity of endometriosis patients who reported CPP [22, 23]. We have shown that TNFα, IL-1β, and IL-6 are elevated in the peritoneal fluids after a single induction of lesions in a mouse model of endometriosis [24, 25]. Lesion induction increases SP, CGRP, and TRPV1 expression in the dorsal root ganglia (DRG) and elevates mechanical hyperalgesia and allodynia [24, 25]. Thus, elevated inflammatory mediators sensitize nociceptor neurons in the endometriotic lesions and/or pelvic organs, initiating pain stimuli, transferring them to the spinal cord and brain to sensitize the central nervous system (CNS), and inducing endometriosis-associated pain.

Immune cells modulate the immune response to inflammation and bi-directionally interact with nociceptors [12]. Macrophages are considered to be key players in promoting endometriosis disease progression and associated pain [27–29], as abundant macrophages are present in ectopic lesions [30] and elevated in the peritoneal cavity [28, 31, 32].

Transcriptionally and functionally dysregulated macrophages can establish an inflammatory environment by secreting cytokines and chemokines that exacerbate innervation and vascularization of lesions [17, 28, 29, 32–34] and contribute to endometriosis-associated pain [32, 35, 36]. Peritoneal macrophages also contribute to the inflammatory condition by releasing cytokines and growth factors that stimulate local inflammation, lesion infiltration, and vascularization [28, 32, 37, 38]. Although peripheral inflammation and sensitization explain some aspects of CPP, CPP can persist or recur in patients after lesion removal [39]. Furthermore, the severity of pain is not correlated with the lesion size, location, and extent of lesion infiltration into tissues [40]. Chronic hyperexcitability perhaps induces long-lasting neuroplastic modification in the CNS.

Neuroinflammation is defined as an inflammatory response within the brain and spinal cord characterized by infiltration of leukocytes, activation of glial cells, and production of proinflammatory cytokines and chemokines [12]. Microglia and astrocytes are key regulators of inflammatory responses within CNS, and the activation of microglial and astrocytes is not only a significant cause of neurologic and neurodegenerative diseases but also painful insults [12, 41]. CPP can also result from CNS top-down activation via neuroinflammation triggered by the dorsal root reflex in the spinal cord to induce peripheral sensitization [12, 42]. In endometriosis, retrograde menstruation, the reflux of menstrual tissues via the fallopian tube into the pelvic cavity, has been widely accepted as the origin of endometriotic lesions [43]. It causes massive inflammatory responses in the peritoneum. However, retrograded menstrual debris is cleared from the pelvic cavity by an innate immune response in the majority of women who do not develop endometriosis [11, 44], but menstrual cycles repeatedly occur in women. Each retrograde menstruation induces composite inflammation in the pelvic cavity, and unsolved inflammation is expected to worsen to develop chronic conditions further [11, 25]. Thus, multiple chronic inflammatory stimuli are expected to enhance central sensitization and induce neuroinflammation, resulting in endometriosis-associated CPP.

In the present study, we carried out repeated cycles of lesion induction to examine how multiple inductions of lesions mimic repeated retrograde menstruation sensitize CNS and whether they can drive neuroinflammation in a mouse model of endometriosis. We also examined mechanical hyperalgesia, peripheral inflammatory mediators and immune cells in the lesions and peritoneal fluids, and neurotransmitters in the DRG to understand how peripheral stimuli are associated with central sensitization and endometriosis-associated pain behavior.

## Materials and Methods

### Animals

C57BL/6 mice were purchased from Inotiv and housed in an environment-controlled animal facility (12:12 light-dark cycle) with ad libitum access to food and water. All animal experiments were performed at Washington State University according to the NIH guidelines for the care and use of laboratory animals (protocol #6751).

### Mouse model of endometriosis

An experimental mouse model of endometriosis was employed by adopting a published procedure with minor modifications [45]. To induce endometriosis-like lesions, female mice (donor) were injected subcutaneously with pregnant mare serum gonadotropin (PMSG, 5 IU, Sigma) to stimulate an estrogenic response within the uterus. Uteri were harvested from donor mice 41 hours after PMSG injection. The endometrium was then separated from the myometrium and dissected into fragments (1-2 mm per side), and 50 mg of fragments were introduced via injection (in 200 µl of PBS) into the peritoneal cavity in the ovary-intact recipient under anesthesia via inhaled isoflurane.

### Study design

Endometriosis-like lesions were induced in the recipient mice for a single time (1x) or six times (6x, at 2-week intervals), as shown in Fig. 1a. On Day –1 (a day before lesion induction), 14, and 42 (2 and 6 weeks after the last induction of 1x or 6x inductions), a behavioral test was performed, and then mice were euthanized for sample collections: peritoneal fluid (PF) was recovered by lavage (4 mL x 2 of ice-cold PBS with 3% FBS), and lesions, bilateral lumbar (L4-6) DRG, spinal cord (L4-6), and brain were collected for further analysis.

**Figure 1.**
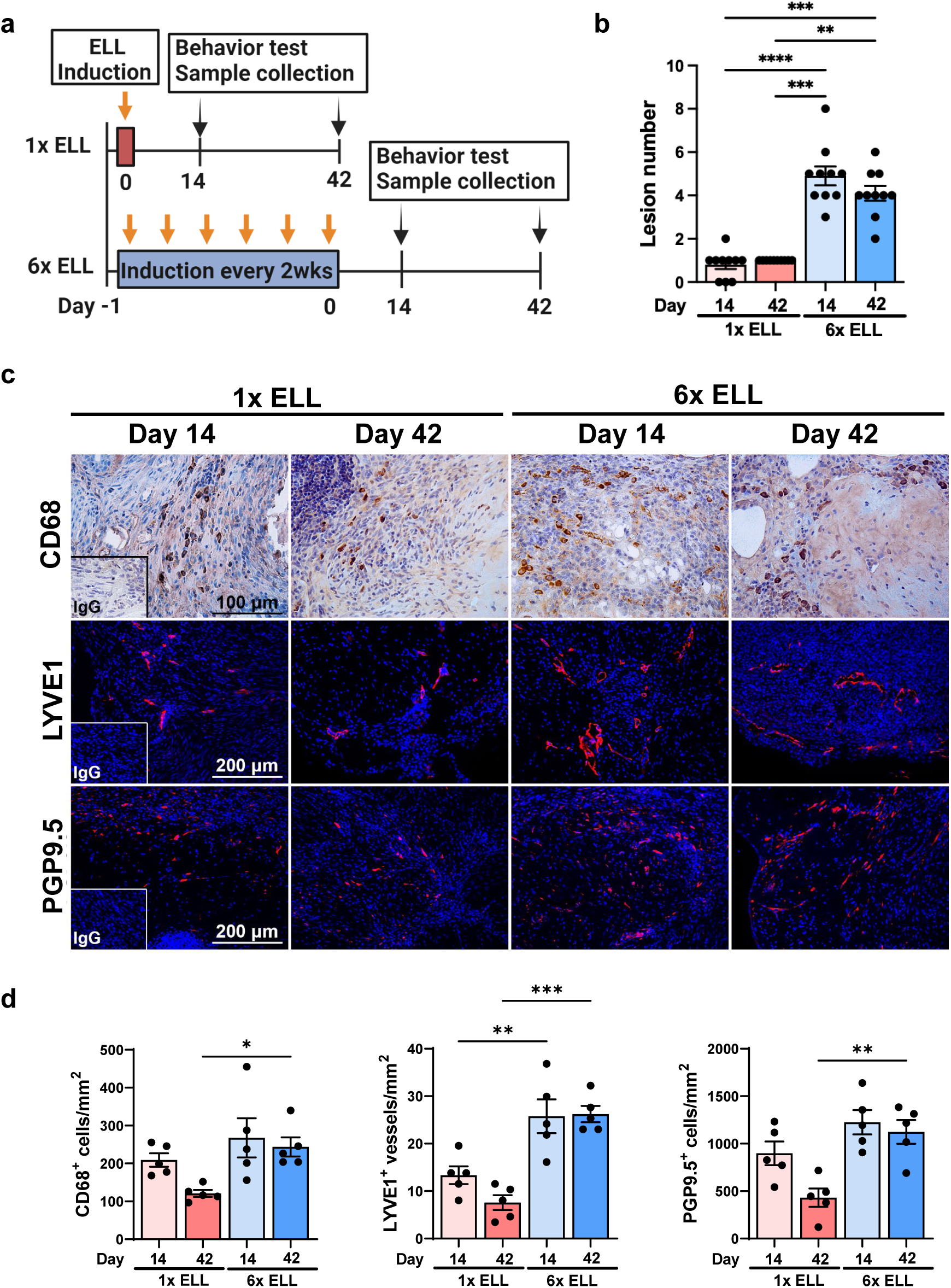
Multiple lesion induction mouse model of endometriosis. (a) Experimental study design as described in Material and Methods. (b) Quantification of lesion numbers in the single or multiple induction mice at 2 or 6 weeks after the last lesion induction (n=10). Representative immunohistochemical images (c) and quantification (d) of CD68+, LYVE1+, or PGP9.5+ cells in the lesions (n=5). Data are shown as the mean ± SEM. ELL: endometriosis-like lesions. \**P* < 0.05, \*\**P* < 0.01, \*\*\**P* < 0.001, \*\*\*\**P* < 0.0001.

### Von Frey test

A standard behavioral (mechanical sensitivity) test was performed before sample collection, as described by our laboratory previously [24, 25]. Mice (n=10/group) were allowed to acclimate in the testing room for 30 min, and then the von Frey test was performed using von Frey filaments (BIO-VF-M, Bioseb). Filaments were applied 10 times to the skin perpendicular to the lower abdomen and bilateral hind paws. The force in grams (g) of the filament evoking a withdrawal response (50% response count as sensitive) was recorded. Three behaviors were considered positive responses to filament stimulation: 1) sharp retraction of the abdomen, 2) immediate licking and/or scratching of the area of filament stimulation, or 3) jumping. All behavioral tests were performed blindly without describing the identity and details of treatment groups to investigators assessing pain. These data were then analyzed by another blinded investigator.

### Flow cytometry

Single-cell suspensions of peritoneal exudate cells were used for analyzing immune cell profiles by flow cytometry as described previously [24, 25, 28, 29]. Briefly, peritoneal exudate cells were lysed using Red Blood Cell Lysis Buffer (BioLegend) and incubated at room temperature for 20 min with Zombie Aqua™ Fixable Viability dye (Bio-Legend). The cells were blocked on ice for 20 min with Fc Block anti-CD16/CD32 (ThermoFisher) and stained with fluorochrome-conjugated monoclonal antibodies for 1 hour (Supplementary Table S1). Samples (n=5/group) were acquired with the Attune NxT Acoustic Focusing Cytometer using Attune NxT software (ThermoFisher), and data were analyzed with FlowJo v10.4 software (FLOWJO).

### IQELISA

Total protein yield from peritoneal fluid was determined by BCA assay (Pierce), and TNFα (IQM-TNFA-1), IL-1β (IQM-IL1b-1), and IL-6 (IQM-IL6-1) were further quantified by IQELISA kits (Ray Biotech) according to the manufacturer’s instructions (n=5/group).

### Immunohistochemistry

Immunostaining of TRPV1, SP, CGRP, PGP 9.5, LYVE1, IBA1, GFAP, neurofilament, and CD68 was performed with cross-sections (5 µm) of paraffin-embedded tissues using specific primary antibodies (Supplementary Table S1) and AlexaFluor 488 or 568-conjugated F(ab’) secondary antibody (Molecular Probe) or VECTASTAIN ABC kit (Vector lab). Immunostaining images were acquired by Leica DM4 B microscopy. Cell-specific CD68-positive cells were counted and quantified by Image J in the area of 0.289768 mm^2^ (n=5/group). LYVE1-positive and PGP9.5-positive cells in the lesion were counted and quantified from three different areas (0.289768 mm^2^/area) using Leica LAS X software (n=5/group). Neurofilament was used as a pan-neuronal marker and was co-stained with TRPV1, SP, or CGRP. TRPV1, SP, or CGRP positive DRG neurons in the section were counted in the area of 0.289768 mm^2^, and the percentages of TRPV1, SP, or CGRP positive cells per neurofilament-positive DRG were shown (n=5/group).

### Image analysis

Image analysis for IBA1 and GFAP was performed as described previously [46] with some modifications. Immunostained IBA1 or GFAP images (1.159063 mm^2^ in size) of the spinal cord (dorsal horn) and the brain (cortex, hippocampus, thalamus, and hypothalamus) were taken and exported by a blinded researcher to avoid any experimental bias. The exported images (1280×960 pixels) were deconvoluted using the inbuilt “Color Deconvolution (H-DAB)” function in Fiji image analysis software to obtain brown-stained areas [47]. The images were loaded into the machine learning “Trainable Weka Segmentation” plugin in Fiji, and the plugin was trained to identify three classes of immunostaining: stained cells, non-stained cells, and background. Then, the images were processed to create a classified image and thresholded [48]. The size and the number of cells were measured using the “Analyze Particles” function in Fiji with a size threshold of 45-infinity. The number of cells was divided by the analyzed area. For determining the percentage area, the total area of immunoreactivity was divided by the analyzed area (n=5/group).

### Statistical analysis

Statistical analyses were performed using GraphPad Prism (version 9.5). Data were tested for normal distribution using the Shapiro-Wilk normality test. If data were normally distributed, one-way ANOVA followed by Tukey multiple comparison tests was used to analyze the differences among the groups. If data were not normally distributed, Mann-Whitney or Kruskal-Wallis test was performed. A P value less than 0.05 was considered to be statistically significant.

## Results

### Endometriosis lesion development and endometriosis-associated hyperalgesia

We first assessed how multiple inoculations of the endometrium affect endometriotic lesion development and progression. Lesion numbers were significantly increased in the multiple induction mice at 2 weeks after the last lesion induction than in mice that received only a single induction (Fig. 1b). These numbers remained higher in the multiple induction mice at 6 weeks after the lesion induction (Fig. 1b). As macrophage infiltration is critical for lesion development, angiogenesis, and innervation [24, 25, 27], we next examined macrophages (CD68), lymphatic endothelial cells (LYVE1), and nerve cells (PGP9.5) in the lesions (Fig. 1cd). CD68+ macrophages were comparable in the single and multiple induction mice at 2 weeks, whereas more CD68+ macrophages were detected in the lesions with multiple inductions at 6 weeks (Fig. 1cd). Abundant LYVE1+ cells were observed in the multiple induction mice compared to the single induction mice at 2 and 6 weeks (Fig. 1cd). Multiple induction mice showed more significant PGP9.5+ nerve cells in the lesions than single induction mice at 6 weeks, although they were not significantly different in the single and multiple induction mice at 2 weeks (Fig. 1cd). Thus, multiple inductions further support endometriotic lesion development and progression by enhancing macrophage infiltration, angiogenesis/lymphangiogenesis, and innervation compared to the single induction. Specifically, macrophage infiltration and innervation remained greater in the multiple induction mice for extended periods.

We next performed the von Frey test to examine the abdominal and hind paw retraction threshold to determine whether multiple lesion inductions affect endometriosis-associated hyperalgesia (Fig. 2). Both single and multiple induction mice withdrew abdominal retraction thresholds with significantly lighter stimuli at 2 and/or 6 weeks than pre-induction mice (Fig. 2a). The multiple inductions showed higher sensitivity than the single induction at 6 weeks (Fig. 2a). The hind paw retraction thresholds were more sensitive in the single and multiple induction mice at 2 weeks than at the pre-induction (Fig. 2b). While the sensitivity of hind paw retraction returned to the pre-induction level at 6 weeks in the single induction mice, it remained high in the multiple induction mice at 6 weeks (Fig. 2b). The results suggest that the multiple induction mice sustain higher sensitivity not only in the abdomen where lesion were established but also a different body site for extended periods, indicating the signs of chronic overlapping pain conditions and/or widespread pain via central sensitization.

**Figure 2.**
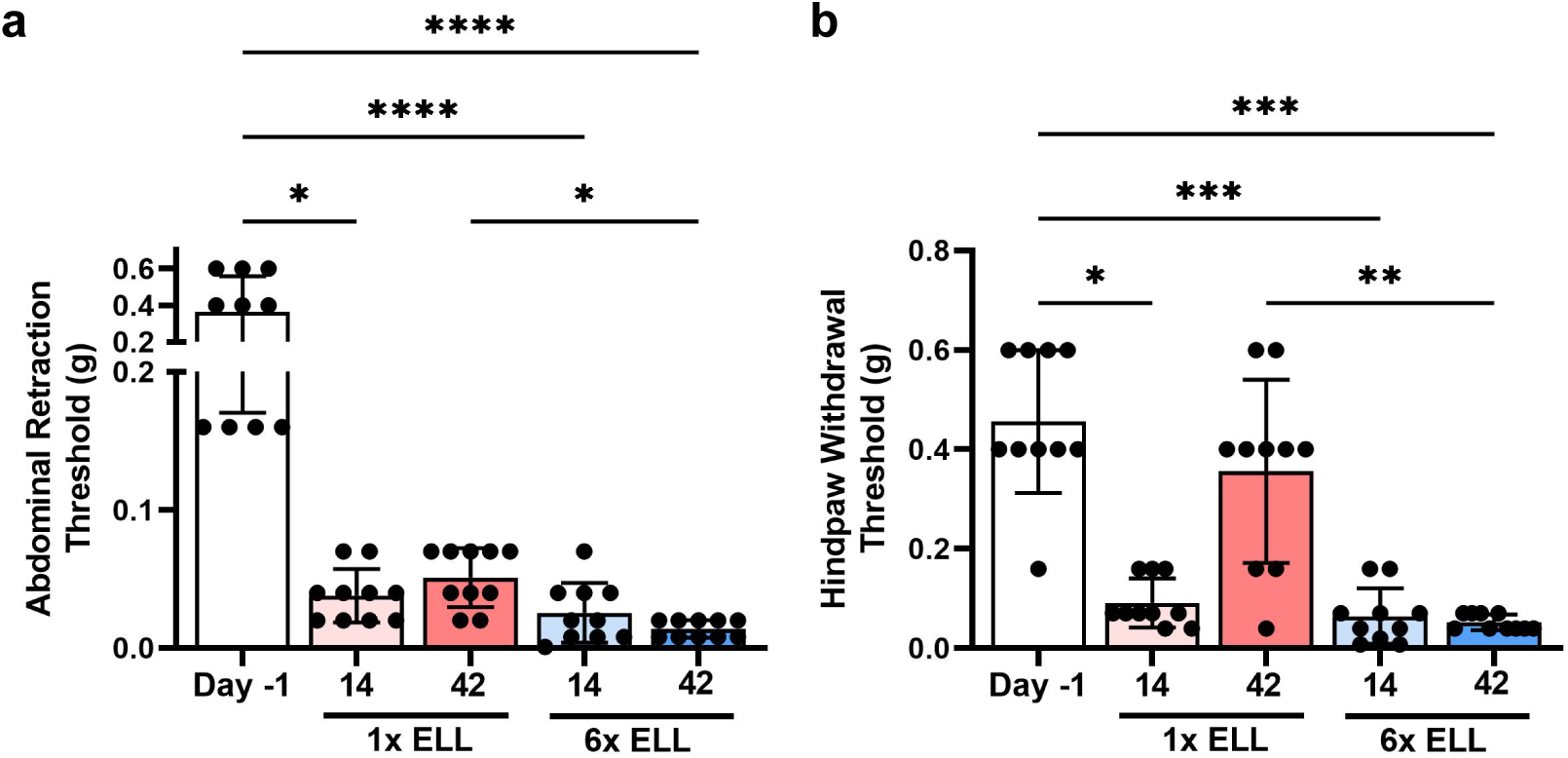
Evaluation of endometriosis-associated hyperalgesia followed by single or multiple inductions at 2 or 6 weeks after the last lesion induction. Abdominal (a) and hind paw (b) withdrawal thresholds were assessed using the von Frey test. Data are shown as mean ± SEM (n = 10). ELL: endometriosis-like lesions. \**P* < 0.05, \*\**P* < 0.01, \*\*\**P* < 0.001, \*\*\*\**P* < 0.0001.

### Microglial activation and astrocytes in the brain and spinal cord

Endometriosis-associated pain can be exacerbated by central sensitization, and glial cells, such as microglia and astrocytes, contribute to developing neuroinflammation and chronic pain [12, 49–51]. Thus, we next analyzed IBA1 (a marker of microglia) and GFAP (a marker of astrocytes) in the brain and spinal cord (Figs. 3-5 and Supplementary Fig. S1). Specifically, the regions of the brain were selected due to the prefrontal cortex for pain processing [52], the hippocampus for pain memory, depression, and anxiety [53, 54], the thalamus for pain modulation and relaying signals [55], and the hypothalamus for mood disorders, stress control, and reproductive function [56].

**Figure 3.**
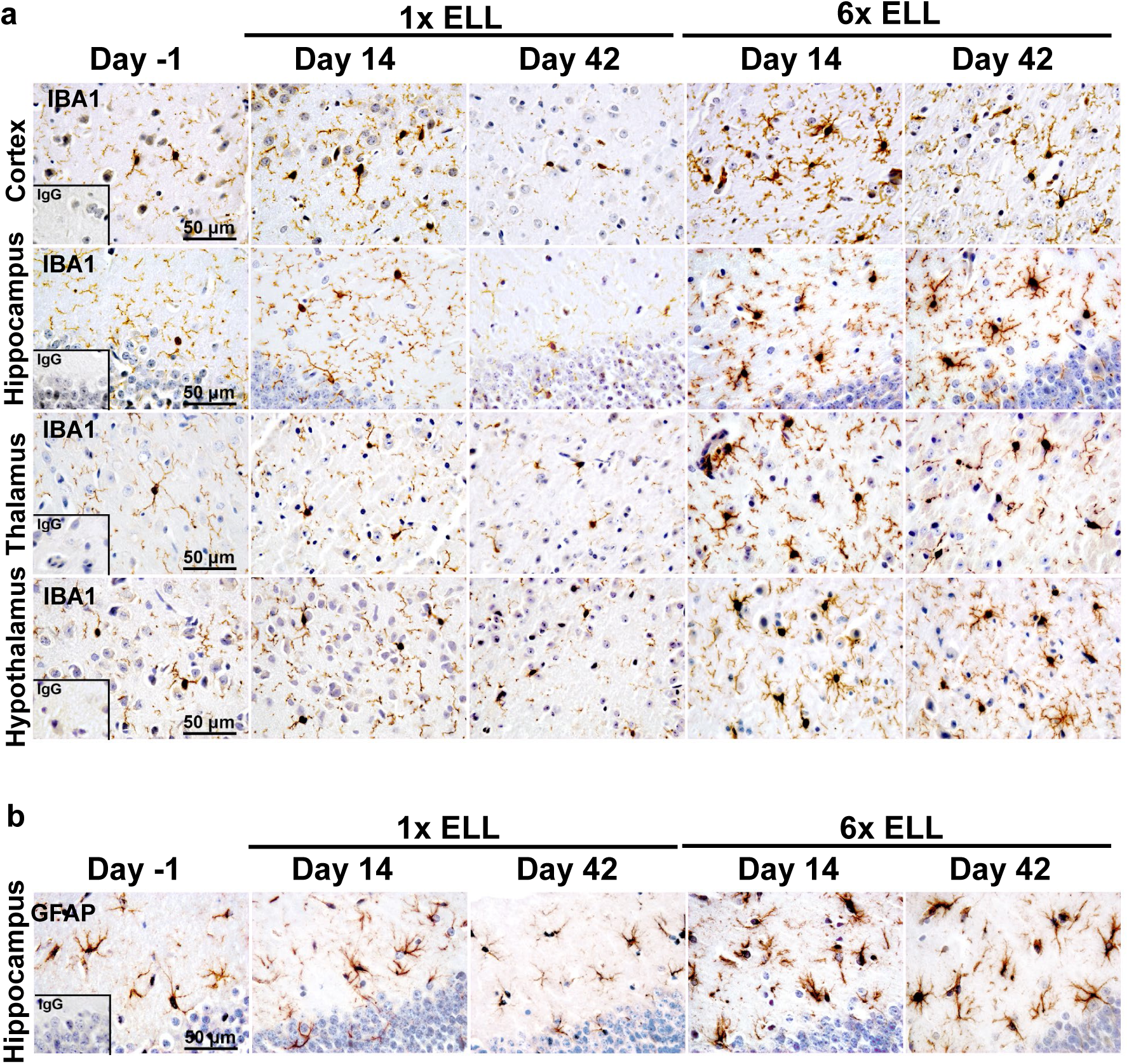
Representative immunohistochemical images of (a) IBA1 in the cortex, hippocampus, thalamus, and hypothalamus, and (b) GFAP in the hippocampus in the single and multiple induction mice at 2 or 6 weeks after the last lesion induction. ELL: endometriosis-like lesions.

As an increase in microglial soma size is considered a key indicator of microglial activation [57, 58], we analyzed the soma size, cell number, and % of cell extended area of IBA1+ microglia, as previously shown [46]. There were no differences in soma size of the microglia within the cortex, hippocampus, thalamus, or hypothalamus of single induction mice at 2 and 6 weeks (Figs. 3a and 4a). In contrast, the microglia of multiple induction mice had significantly enlarged somas in the hippocampus at 2 and 6 weeks and in the thalamus at 2 weeks compared with those in pre-induction mice (Figs. 3a and 4a). Soma size in the hippocampus or thalamus of multiple induction mice at 6 weeks or 2 and 6 weeks, respectively, was greater than that of single induction mice at these same time points (Figs. 3a and 4a). IBA1+ microglia number and/or % of area were increased in the hippocampus and/or hypothalamus of single induction mice only at 2 weeks. However, they were elevated in the cortex, hippocampus, thalamus, and hypothalamus of multiple induction mice at both 2 and 6 weeks (Figs. 3a and 4a). Furthermore, multiple inductions induced more IBA1+ microglia number or % of area in most brain regions than single induction, some at 2 weeks but all at 6 weeks (Figs. 3a and 4a).

**Figure 4.**
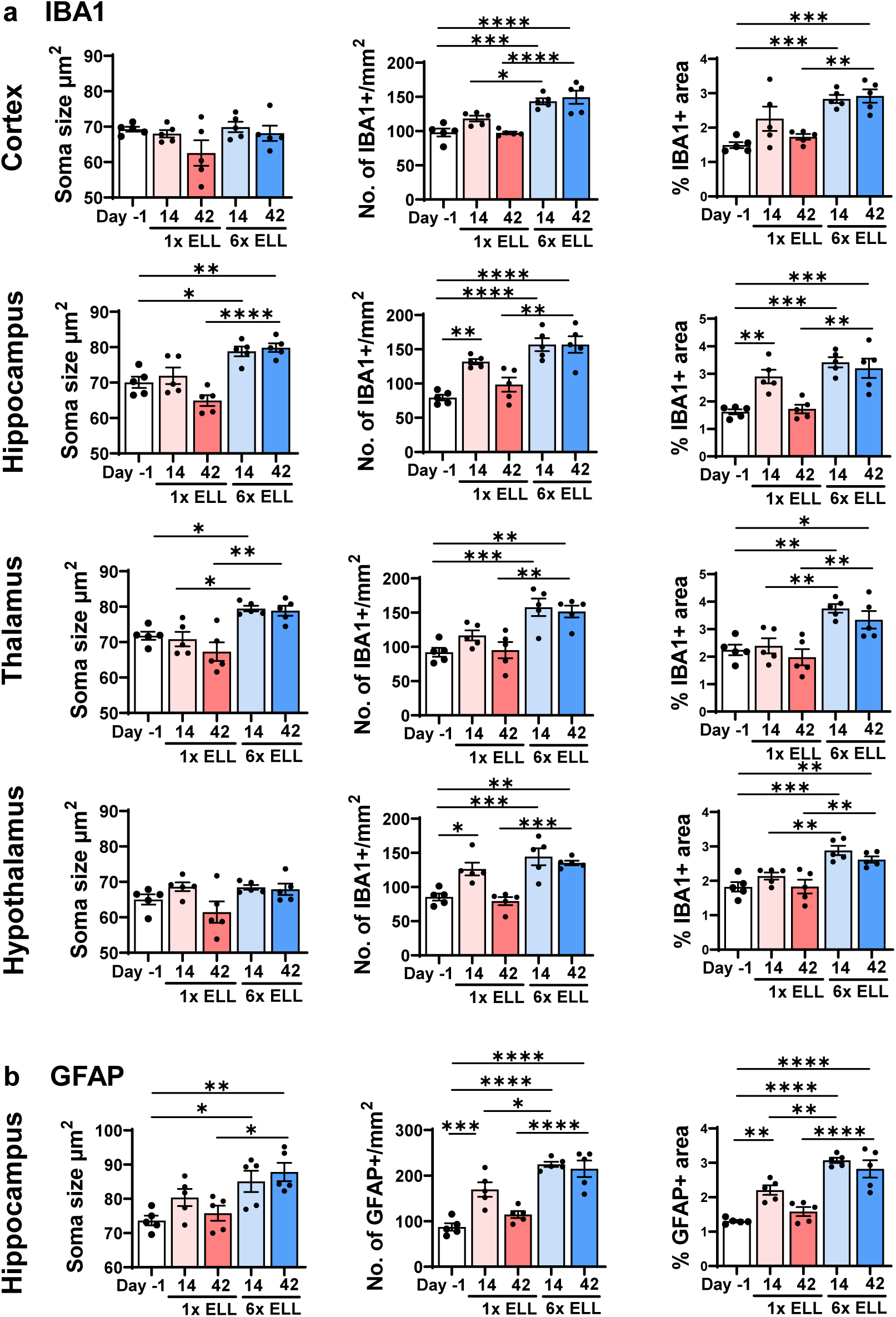
Quantification of immunohistochemical images of (a) IBA1 in the cortex, hippocampus, thalamus, and hypothalamus, and (b) GFAP in the hippocampus in the single and multiple induction mice at 2 or 6 weeks after the last lesion induction. Data are shown as mean ± SEM (n = 5). ELL: endometriosis-like lesions. \**P* < 0.05, \*\**P* < 0.01, \*\*\**P* < 0.001, \*\*\*\**P* < 0.0001.

**Figure 5.**
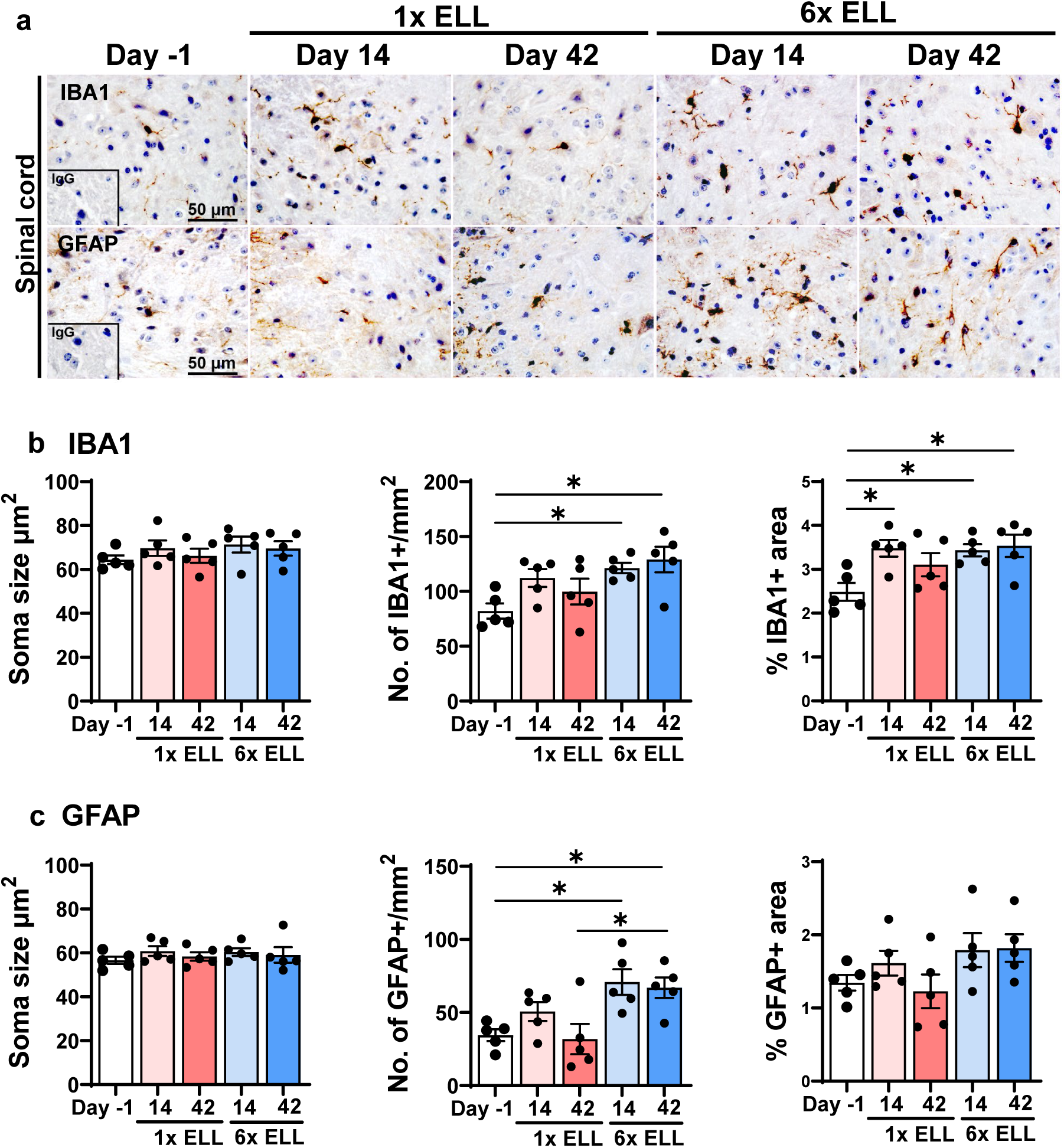
Representative immunohistochemical images (a) and quantification (bc) of IBA and GFAP in the spinal cord in the single and multiple induction mice at 2 or 6 weeks after the last lesion induction. Data are shown as the mean ± SEM (n=5). ELL: endometriosis-like lesions. \**P* < 0.05.

Astrocyte-mediated neuroinflammation is also a key mechanism underlying the maintenance of chronic pain [12, 59, 60]. Chronic neuropathic pain is known to induce astrocyte swelling [61]. Thus, we next analyzed astrocytes in the brain regions (Figs. 3b and 4b, and Supplementary Fig. S1ab), following the evaluation methods of microglia. In the hippocampus, the soma size of the astrocytes was larger in the multiple induction mice than in pre-induction mice at 2 and 6 weeks, but unchanged in the single induction mice (Figs. 3b and 4b). At 6 weeks, the soma size of the astrocytes was greater in the multiple induction mice than in the single induction mice (Figs. 3b and 4b). GFAP+ astrocyte number and % of area were elevated in the single induction mice at 2 weeks and in the multiple induction mice at 2 and 6 weeks compared with those at pre-induction. Multiple inductions further increased GFAP+ astrocyte number and % than single induction at both time points (Figs. 3b and 4b). In contrast, the soma size of the astrocytes did not alter in the cortex, thalamus, and hypothalamus following single or multiple lesion inductions (Supplementary Fig. S1ab). GFAP+ astrocyte number and % of area were elevated in the hypothalamus of multiple induction mice at 2 and 6 weeks, and % of GFAP+ area was higher in the cortex (Supplementary Fig. S1ab).

In the spinal cord, the soma size of microglia and astrocytes was not altered by lesion induction (Figs. 5ab). Multiple inductions induced more IBA1+ microglia number and % of area compared with those in pre-induction mice, whereas single induction only increased % of IBA1+ area at 2 weeks (Figs. 5ab). GFAP+ astrocyte number was also elevated in the spinal cord by multiple inductions at 2 and 6 weeks, and the number was higher in the multiple induction mice than in the single induction mice at 6 weeks (Figs. 5ab).

### Pain-related mediators in the DRG

DRG are sensory neurons that detect and transmit stimuli to the CNS [62]. We have reported increased expression of transient receptor potential channels, TRPV1, and neurotransmitters, such as SP and CGRP, in mouse endometriosis [25]. We thus examined TRPV1, SP, and CGRP in the L4-6 DRG, the primary spinal ganglia receiving sensory input from pelvic organs (Fig. 6). Both single and multiple lesion inductions increased TRPV1, SP, and CGRP expression at 2 weeks compared with those at pre-induction (Fig. 6ab). Elevated TRPV1+ and SP+ DRG remained high in the multiple induction mice at 6 weeks but not in the single induction mice, while CGRP+ DRG were still high in the single induction mice at 6 weeks (Fig. 6ab). Furthermore, more SP+ and CGRP+ DRG were detected in the multiple induction mice than in the single induction mice at 2 and 6 weeks (Fig. 6ab). These results indicate that multiple inductions induce prolonged stimulation of nociceptor neurons in the DRG.

**Figure 6.**
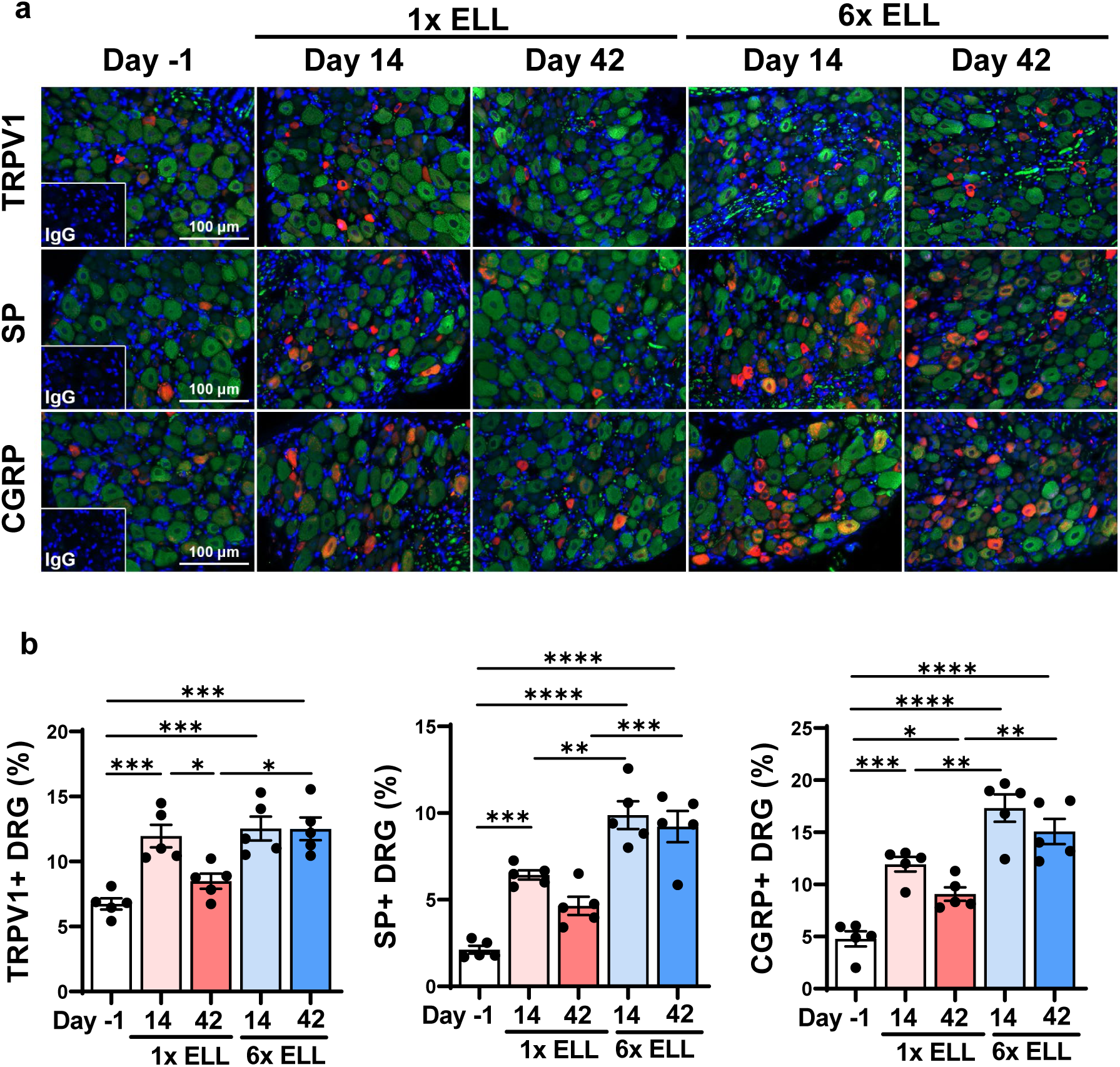
Expression of TRPV1, SP, and CGRP in DRG in the single and multiple induction mice at 2 or 6 weeks after the last lesion induction. (a) Representative images showing DRG sections double stained with TRPV1, SP, or CGRP (red), and neurofilament (green), as a marker of neural cells. (b) Quantification of TRPV1+, SP+, or CGRP+ cells in neurofilament-positive cells. Data are shown as the mean ± SEM (n=5). ELL: endometriosis-like lesions. \**P* < 0.05, \*\**P* < 0.01, \*\*\**P* < 0.001, \*\*\*\**P* < 0.0001.

### Peritoneal macrophage dynamics and inflammatory environment establishment in the peritoneal cavity

Heterogenous macrophage populations time-dependly alter in the peritoneum after lesion induction in mice [25]. We next examined how multiple inductions affect proinflammatory macrophages (TIM4^lo^ MHCII^hi^), FRβ+ macrophages, and residential macrophages (TIM4^hi^ MHCII^lo^), as well as neutrophils (Ly6G+) (Fig. 7). Although there were no significant differences in the CD11b+ total macrophage population between single and multiple inductions at 2 and 6 weeks, Ly6G+ neutrophils were significantly elevated in the multiple induction mice at 2 weeks (Fig. 7ad). CD11b+ macrophages were further gated to TIM4^lo^ MHCII^hi^ and TIM4^hi^ MHCII^lo^ macrophages to examine proinflammatory and residential macrophages, respectively (Fig. 7b). Both single and multiple inductions reduced TIM4^hi^ MHCII^lo^ macrophages at 2 weeks as a sign of macrophage disappearance reaction (MDR). The population of TIM4^hi^ MHCII^lo^ macrophages at 2 weeks was lower in the multiple induction mice than in the single induction mice (Fig. 7be), suggesting that the multiple inductions induced severe MDR. At 6 weeks, residential macrophages in the single induction mice returned to the pre-induction level but were still lower in the multiple induction mice. Thus, the MDR induced by the single induction was replenished and recovered, but the MDR induced by multiple inductions was not entirely resolved at 6 weeks (Fig. 7be). The single and multiple inductions elevated TIM4^lo^ MHCII^hi^ proinflammatory macrophages at 2 weeks, while the multiple inductions further elevated their populations (Fig. 7be). TIM4^lo^ MHCII^hi^ macrophages returned to the pre-induction levels in both groups at 6 weeks (Fig. 7be). We have previously reported the FRβ+ macrophage population that was differentiated from monocyte-derived proinflammatory macrophages and possessed residential macrophage characteristics [29]. The single and multiple inductions elevated FRβ+ macrophages at 2 weeks compared to those in pre-induction level (Fig. 7cf). FRβ+ macrophages were higher in the multiple induction mice than in the single induction mice at 2 weeks (Fig. 7cf). High levels of FRβ+ macrophages were sustained at 6 weeks in the multiple induction mice (Fig. 7cf). When FRβ+ macrophages were further gated to TIM4+ or MHCII^hi^, most of the FRβ+ macrophages expressed high MHCII but limited TIM4 expression after lesion induction (Fig. 7cf). Specifically, MHCII^hi^ FRβ+ macrophages were significantly elevated by the multiple inductions at 2 weeks (Fig. 7cf). These results suggest that elevated FRβ+ macrophages after lesion inductions were newly recruited monocyte-derived highly inflammatory macrophages, and the multiple inductions further recruited and elevated them in the peritoneal cavity.

**Figure 7.**
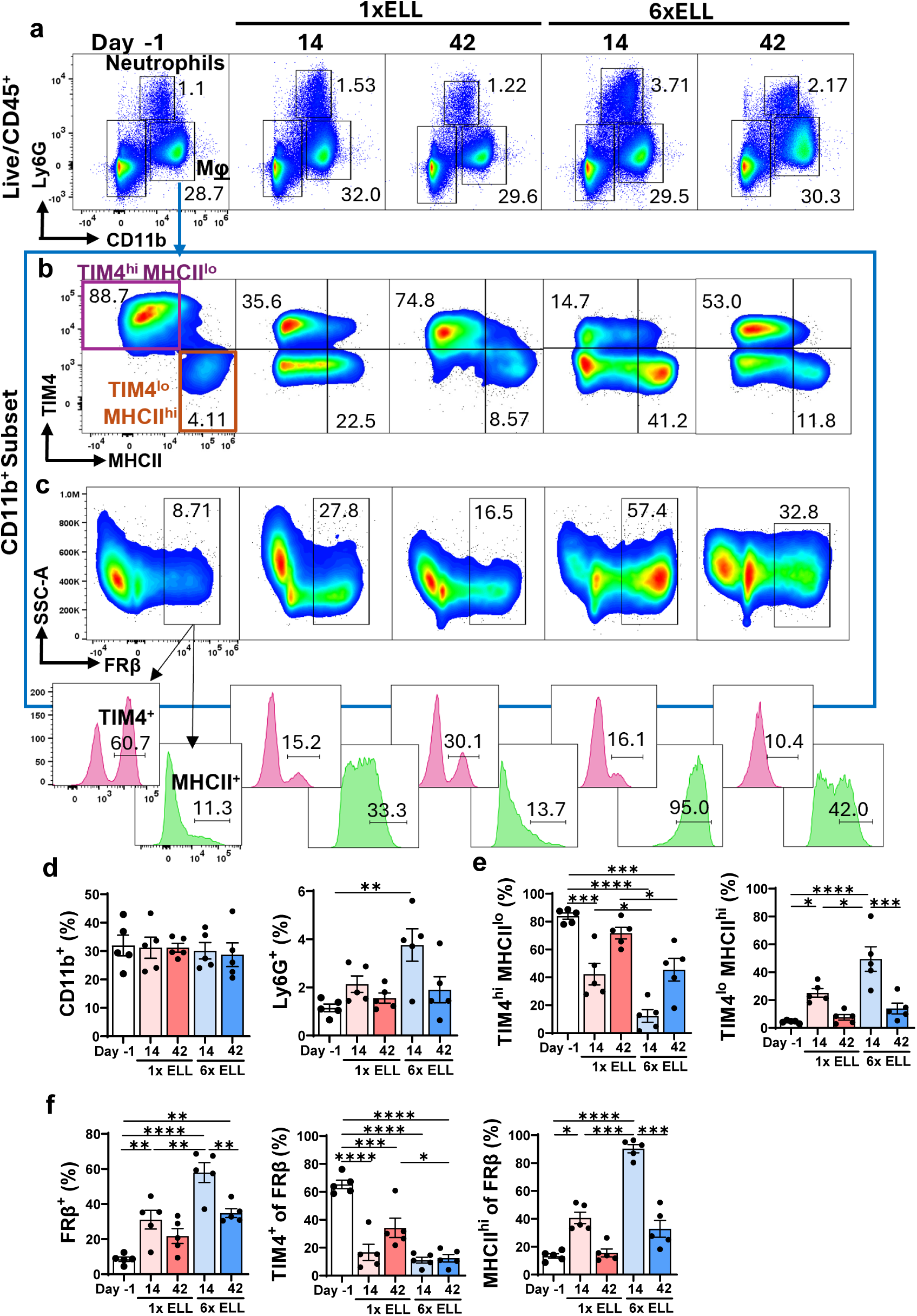
Comparison of peritoneal immune cell profiles in the single and multiple induction mice at 2 or 6 weeks after the last lesion induction. (a) Representative flow plots illustrating the composition of CD11b+ and Ly6G+ cells. (b) CD11b+ cells were further gated by TIM4 and MHCII. (c) CD11b+ cells were further gated by FRβ (top), and FRβ+ cells were then gated by TIM4 and MHCII (bottom). Proportions of CD11b+ or Ly6G+ (d) and TIM4^hi^ MHCII^lo^ and TIM4^lo^ MHCII^hi^ (e) are shown. (f) Proportions of FRβ+ of CD11b+ cells, and TIM4+ or MHCII^hi^ of FRβ+ macrophages were shown. Data are shown as the mean ± SEM (n=5). ELL: endometriosis-like lesions. \**P* < 0.05, \*\**P* < 0.01, \*\*\**P* < 0.001, \*\*\*\**P* < 0.0001.

In addition to macrophages, we also examined peritoneal B– and T-cells (Supplementary Fig. S2). CD19+ B cells were reduced in the multiple induction mice at 2 weeks compared with those in the pre-induction mice (Supplementary Fig. S2ac). CD3+ T-cells were elevated at 2 weeks in the multiple induction mice following increased CD8+ and CD4+ T-cells (Supplementary Fig. S2abd). CD4+ T-cells were higher at 6 weeks in the multiple induction mice than the single induction mice (Supplementary Fig. S2bd).

To confirm elevated inflammation via the multiple inductions, peritoneal TNFα, IL-1β, and IL-6 protein concentrations were assessed (Fig. 8), as these cytokines are considered the key factors involved in maintaining the aberrant peritoneal inflammatory environment, promoting lesion growth and mediating peripheral sensitization [63–65]. The single and multiple inductions significantly elevated secreted TNFα, IL-1β, and IL-6 levels in the peritoneal cavity (Fig. 8) at 2 weeks. All cytokine levels were higher in the multiple induction mice than in the single induction mice at 2 weeks (Fig. 8). Furthermore, elevated cytokine levels returned to the pre-induction levels in the single induction mice at 6 weeks, however, they remained high in the multiple induction mice (Fig. 8). These results further support that the multiple inductions establish the aberrant inflammatory environment in the peritoneal cavity.

**Figure 8.**
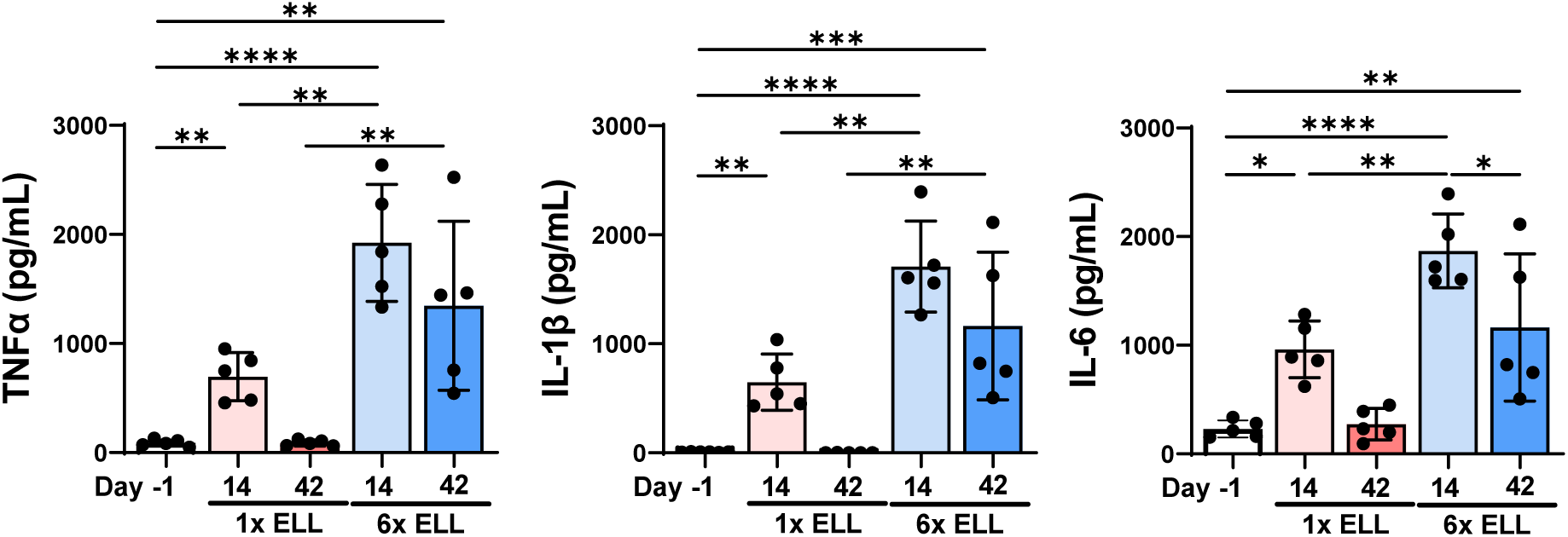
Proinflammatory cytokine levels (TNFα, IL-1β, and IL-6) in the peritoneal fluid were analyzed by IQELISA. Data are shown as the mean ± SEM (n=5). ELL: endometriosis-like lesions. \**P* < 0.05, \*\**P* < 0.01, \*\*\**P* < 0.001, \*\*\*\**P* < 0.0001.

## Discussion

Approximately 60-80% of women with endometriosis suffer endometriosis-associated CPP [66, 67], which is 13 times higher than healthy patients [67]. Endometriosis patients experience menstrual cyclic and acyclic pain, i.e. dysmenorrhea with dyschezia, dysuria, or dyspareunia [66], and pain can be expanded throughout the pelvis and abdomen, further referred to the back and legs [66]. Women with endometriosis are often diagnosed with bladder and colon sensory dysfunctions, such as irritable bowel syndrome (IBS) and/or overactive bladder syndrome (OAB) [68]. Widespread pain is also a common experience in women with endometriosis. Phan et al. [69] have reported that endometriosis-associated CPP often causes myofascial dysfunction and sensitization beyond the pelvic regions that may be initiated or maintained by ongoing pelvic floor spasms. These comorbidities indicate widely varied endometriosis-associated CPP and more complex pathophysiology of endometriosis. Recent evidence suggests that protracted peripheral and central sensitization are present in endometriosis patients with CPP [11]. In the present study, we designed to induce multiple endometrial inoculations to mimic retrograde menstruation, as mice do not have menstrual cycles. As an important phenotype, our study demonstrated that multiple inductions of lesions resulted in greater hyperalgesia, especially presenting increased prolonged hind paw sensitivities in addition to abdominal sensitivity. While abdominal sensitivity is considered peripheral visceral pain due to thinner skin and less underlying muscle, the hind paw can be affected by both peripheral and central sensitization processing neural pathways [70]. Although lesion numbers were increased by multiple inductions as a nature of the mouse model of endometriosis (>90% of mice develop lesions, which could be a limitation of the study), endometriosis-associated pain is not correlated with disease extent in women with endometriosis [11]. Thus, endometriotic lesion-dependent pain is apparent; however, these lesions cannot be the sole source of endometriosis-associated CPP.

Our results showed prolonged glial activation in several brain regions in the multiple induction mice. A consistent increase in the soma size of microglia and/or IBA+ microglial cells was observed in the brain and spinal cord, which indicates characteristic features of neuroinflammation in the CNS. Interestingly, the larger soma size of microglia and astrocytes with elevated IBA+ or GFAP+ cells was only observed in the hippocampus. Many studies have reported hippocampus abnormalities in patients experiencing chronic pain, anxiety, and depression [71]. GFAP+ astrocytes in the hippocampus are associated with mood disorders in persistent pain states [60, 71]. Endometriosis is known to affect the mental health and emotional well-being of women, leading to anxiety and depression [72, 73]. Due to abundant glial activation in the hippocampus induced by multiple inductions, cyclic sources of peripheral input are likely to induce neuroinflammation for extended periods, causing anxiety and depression and reducing the quality of life in endometriosis women. IBA1+ microglial cells were increased in the cortex, which has important pain-processing functions connecting stimuli to other brain regions, such as the hippocampus and thalamus [52]. As-Sanie et al. [74, 75] demonstrate that changes in regional gray matter volume within the central pain system in the cortex play an important role in developing endometriosis-associated CPP, regardless of the endometriotic lesions. While the connection between neuroinflammation and the altered gray matter volume in the cortex is unclear, the changes in the central pain system are crucial to developing endometriosis-associated CPP. In addition to the hippocampus and/or cortex, we have observed a persistent increase of IBA1+ and GFAP+ cells in the hypothalamus in the multiple induction mice. Microglia in the hypothalamus are considered to be key regulators of homeostasis processes, transmitting sensing signals to the CNS [76]. Microglia can regulate the hypothalamus-pituitary-adrenal (HPA) axis with the involvement of the stress process in controlling cortisol levels [77, 78]. Neuroinflammation in the hypothalamus can also alter the HGA axis and develop glucocorticoid resistance associated with somatic diseases and depressive disorders [79]. Thus, our results support the contribution of hypothalamus neuroinflammation for endometriosis-associated anxiety and depression.

Increased soma size of microglia has been reported in the cortex, hippocampus, thalamus, and hypothalamus in a mouse model of endometriosis with a single induction of lesion [46]. In contrast, single lesion induction in our study did not show strong glial activation, except IBA1+ microglia and GFAP+ astrocytes in the hippocampus or hypothalamus or IBA1+ microglia in the spinal cord at 2 weeks. However, it should be noted that a different method was used to induce lesions in the previous study [46]. Chiefly, the uterine fragments were inoculated by a dorsolateral incision [46], whereas we chose to inject minced endometrial tissues with a needle to reduce the amount of procedural-specific inflammatory stimulation. We thus assume that the higher stimuli were induced by the cutting and suturing of the skin and muscle layer than the simple injection. In support of this, ovariectomy “surgery” can increase macrophage replenishment and alter the peritoneal immune environment [80].

In the present study, multiple lesion inductions elevated peripheral inflammation due to high and persistent TNFα, IL-1β, and IL-6 levels in the peritoneal fluid for extended periods. In contrast, single induction only increased cytokine levels up to 2 weeks after lesion induction, meaning initial inflammation has probably been resolved. The results of immune cell distribution in the peritoneal cavity support establishing a chronic inflammatory environment via multiple inductions. Peritoneal macrophages are highly diverse [29, 80], differ in their ontogeny [81], and have transcriptionally and functionally divergent features depending on the signals of the local environment [82]. When endometrial tissues are introduced in the peritoneum, acute inflammatory responses are caused. Peritoneal residential macrophages (TIM4^hi^ MCHII^lo^) are important for the initial uptake where they adhere to the mesothelium to cover organs [83, 84] or die via pyroptosis to release proinflammatory cytokines, such as IL-1β [85], called MDR. If residential macrophages die/disappear, they appear to be replaced by bone marrow/monocyte-derived macrophages [86]. Our study showed that MDR induced by multiple inductions was more severe than that in the single induction. In support of our previous study [25], MDR was recovered by 6 weeks in the single induction mice, whereas MDR was not fully solved at 6 weeks in the multiple induction mice. Following MDR results, a more significant monocyte-derived proinflammatory macrophage population was found in the multiple induction mice, indicating higher levels of inflammation with severe replenishment of macrophages have occurred. Interestingly, Ly6G+ neutrophils were also elevated in the multiple induction mice at 2 weeks. Neutrophils are first to arrive in the peritoneal cavity when inflammation occurs as an initial inflammatory response and die immediately after [87]. Thus, persistent inflammatory stimuli still exist in the peritoneal cavity 2 weeks after lesion induction in the multiple induction mice. Our previous study demonstrates that monocyte-derived proinflammatory macrophages further differentiate into FRβ+ macrophages with some residential macrophage features (=large peritoneal macrophages) [29]. Herein, we show that newly recruited FRβ+ macrophages highly express MHCII but lowly express TIM4. These results suggest that repetitive inoculations of endometrial tissues cause persistent inflammatory stimuli to enhance and maintain peripheral chronic inflammation, probably elevating FRβ+ macrophages. Because neurotransmitters (SP and CGRP) and TRPV1 were greater in the DRG in the multiple induction mice, chronic inflammatory stimuli further affect the peripheral sensory nervous system. Of note, the peritoneal T-cell population was increased in multiple induction mice, which was not seen in our previous study using a single induction mouse endometriosis [24, 25, 28]. CD8+ T cells have been reported to be enriched in the endometriotic lesions, potentially linked to endometriosis development, infertility, and chronic pain [88, 89]. Further involvement of T-cell functions and CPP remains to be studied.

In the present study, we used a multiple induction mouse model of endometriosis to mimic repeatedly occurring retrograde menstruation to study how endometriosis-associated CPP has been established. We demonstrate that multiple inductions can enhance peripheral sensitization via established chronic inflammation with altered peritoneal macrophage profiles. We have also found that multiple inductions of lesions induce persistent glial cell activation as a sign of neuroinflammation across several brain regions linked to pain processing, anxiety, depression, and stress response. Neuroinflammation can give feedback to stimulate peripheral organs, potentially inducing widespread pain in endometriosis patients. Indeed, the multiple induction mice showed higher endometriosis-associated hyperalgesia than the single induction mice. Especially hind paw sensitivity was persistent in the multiple induction mice, although anxiety and depression-related behavioral tests should be included in future studies. Thus, repeatedly occurring retrograde menstruation can be the peripheral stimuli that induce nociceptive pain but also induce composite chronic inflammatory stimuli, which may cause neuroinflammation and further sensitize CNS. The circuits of neuroplasticity from enhanced chronic inflammation and stimulation of peripheral organs via the feedback loop of neuroinflammation may induce widespread endometriosis-associated CPP. It is known that the presence of endometriosis lesions does not appropriately explain endometriosis-associated CPP, and additional mechanisms to understand dysfunctions in the CNS can be crucial [66, 74, 75, 90, 91]. While many studies focus on lesion formation and development in the pathogenesis of endometriosis, it will be necessary to study underlying mechanisms for the endometriosis-associated CPP to understand endometriosis pathophysiology further.

## Author contributions

M.S. and K.H. designed the research; M.H., M.S., Y.O., and D.M. performed research and analyzed data; J.A.M. and O.D.S. provided critical feedback on the manuscript; K.H. wrote the paper; all authors read, reviewed, edited, and approved the manuscript.

## Availability of data and materials

The raw data and images used and analyzed for the study are available upon reasonable request.

## Ethics approval and consent to participate

All animal experiments were performed at Washington State University according to the NIH guidelines for the care and use of laboratory animals (protocol #6751).

## Consent for publication

Not applicable.

---

**Supplementary Figure S1.** Representative immunohistochemical images (a) and quantification (b) of GFAP in the cortex, thalamus, and hypothalamus. Data are shown as the mean ± SEM (n=5). ELL: endometriosis-like lesions. \*\**P* < 0.01, \*\*\**P* < 0.001, \*\*\*\**P* < 0.0001.

**Supplementary Figure S2.** Comparison of peritoneal B or T cell profiles in the single and multiple induction mice at 2 or 6 weeks after the last lesion induction. (a) Representative flow plots illustrating the composition of CD19+ and CD3+ cells. (b) CD3+ cells were further gated by CD8 and CD4. Proportions of CD19+ or CD3+ (c) and CD8+ or CD4+ (d) are shown. Data are shown as the mean ± SEM (n=5). ELL: endometriosis-like lesions. \**P* < 0.05, \*\**P* < 0.01, \*\*\*\**P* < 0.0001.

