## Supplementary figures and images for "Multiple lesion inductions intensify central sensitization driven by neuroinflammation in a mouse model of endometriosis"

### Suppl Figures

#### Supplementary Figure 1

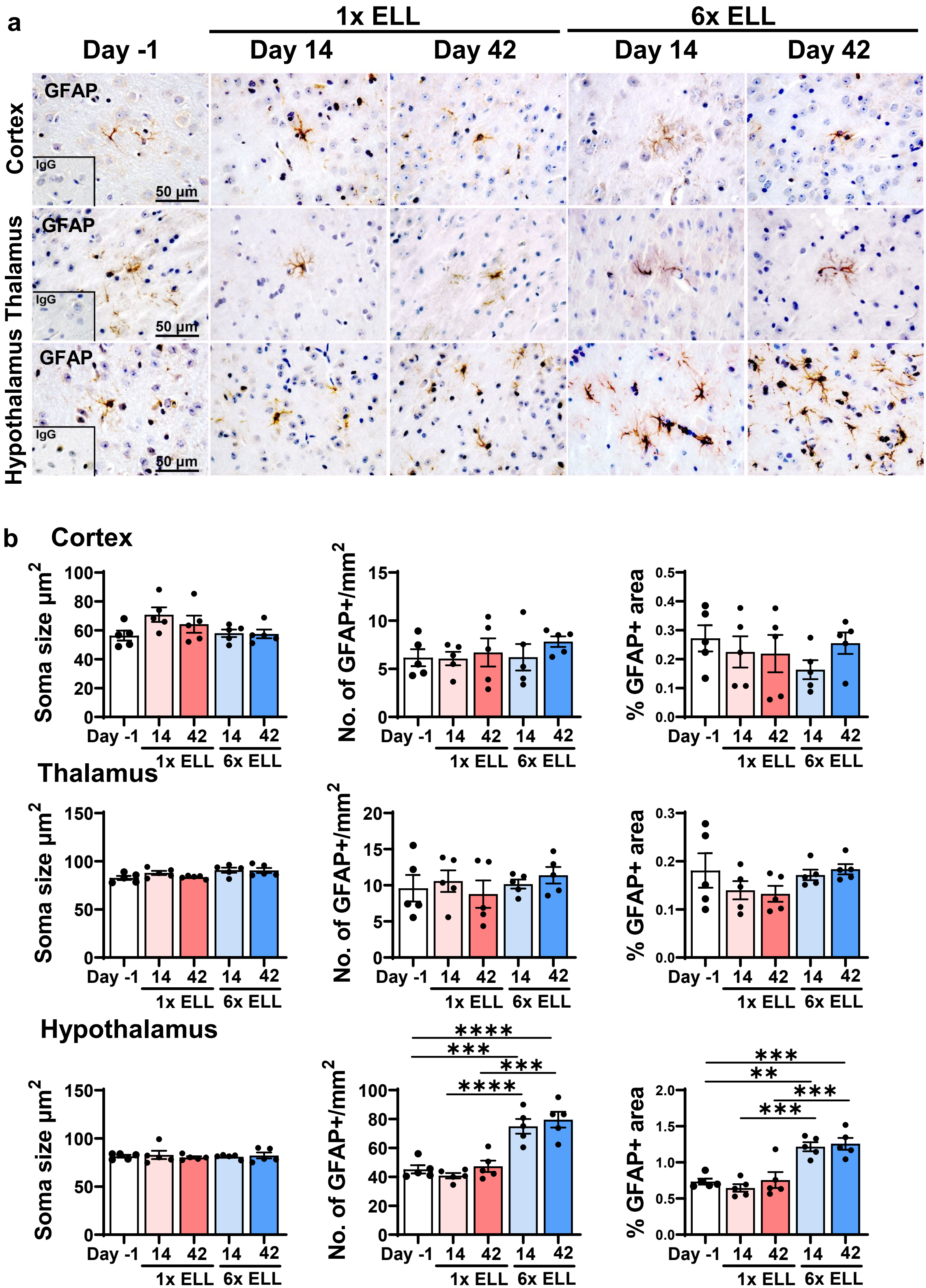

#### Supplementary Figure 2

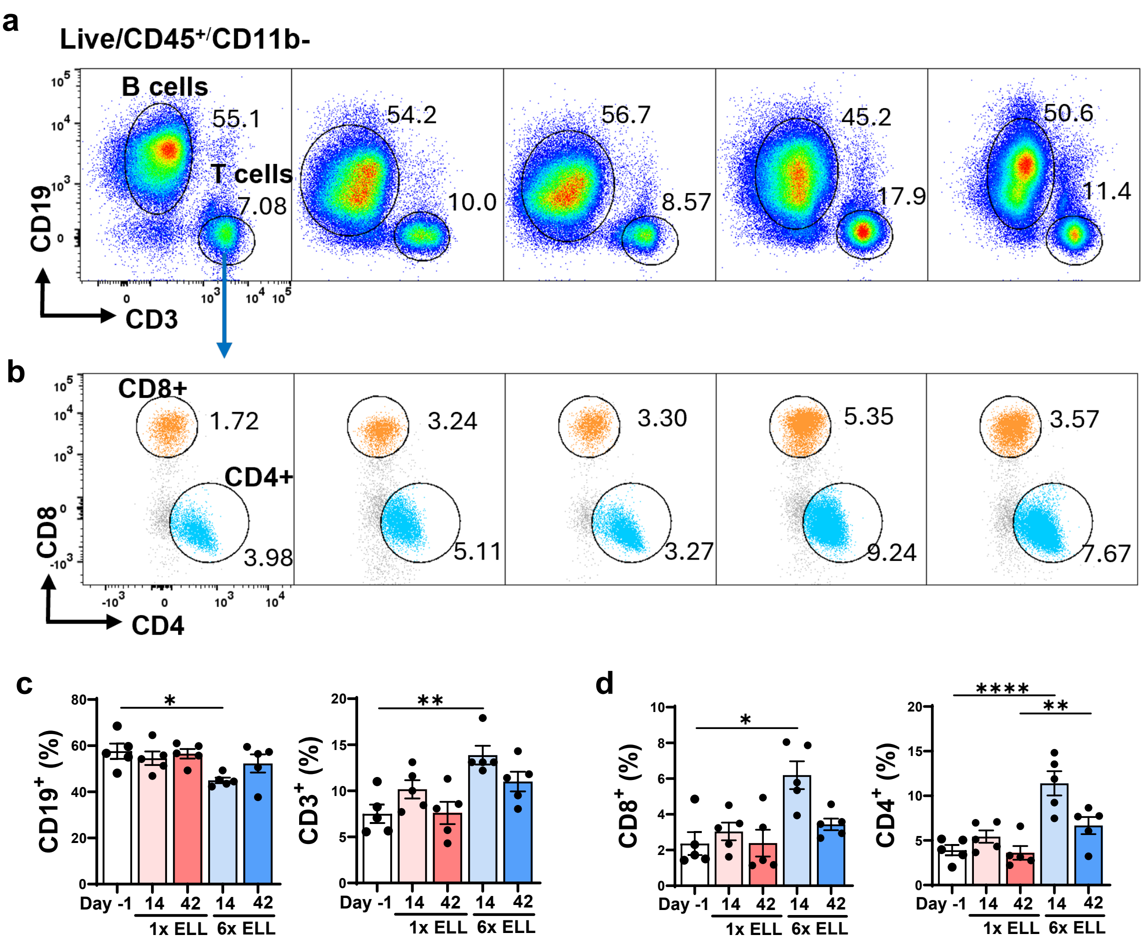
