## Supplementary material for "Multiple lesion inductions intensify central sensitization driven by neuroinflammation in a mouse model of endometriosis": Suppl Table

### Supplementary Table S1 Antibodies and Reagents for Flow Cytometry and Immunochemistry

| Antibody | Conjugate | Company | Catalog# | Application |
|---|---|---|---|---|
| CD11b (M1/70) | APC-Cy7 | BD Biosciences | 557657 | Flow |
| CD19 (ID3) | NovaFluor Yellow 610 | eBioscience™ | M004T02Y03 | Flow |
| CD3 (17A2) | FITC | BioLegend | 100203 | Flow |
| CD45 (30-f11) | PE-Cy5 | BioLegend | 103109 | Flow |
| CD68 (KP1) | Unconjugated | Abcam | ab955 | IHC |
| CGRP (4901) | Unconjugated | Abcam | ab81887 | IHC |
| GFAP | Unconjugated | Abcam | ab7260 | IHC |
| IA/IE, MHC II (M5/114) | BV711 | BD Biosciences | 563414 | Flow |
| IBA1 | Unconjugated | Wako | 019-19741 | IHC |
| Ly6G (1A8) | Super Bright 436 | eBioscience™ | 62-9668-82 | Flow |
| LYVE1 | Unconjugated | AngioBio | 11-034 | IHC |
| Neurofilament | Unconjugated | Millipore | AB5539 | IHC |
| PGP9.5 | Unconjugated | Invitrogen | PA5-29012 | IHC |
| Substance P (SP-DE4-21) | Unconjugated | Abcam | ab14184 | IHC |
| TIM4 (RMT4-54) | PE | BioLegend | 130005 | Flow |
| TRPV1 (VR1-C) | Unconjugated | Neuromics | RA14113 | IHC |
| Fc Block CD16/CD32 antibody | | Thermo Fisher | 14-0161-82 | Flow |
| Total Antibody Compensation Bead Kit | | Thermo Fisher | A10513 | Flow |
| Zombie Aqua™ Fixable Viability Kit | | BioLegend | 423101 | Flow |
